# Antibody-free time-resolved terbium luminescence assays designed for cyclin-dependent kinase 5 (CDK5)

**DOI:** 10.1101/2024.04.24.590988

**Authors:** Jason L. Heier, Dylan J. Boselli, Laurie L. Parker

## Abstract

Novel time-resolved terbium luminescence assays were developed for CDK5 and CDK2 by designing synthetic substrates which incorporate phospho-inducible terbium sensitizing motifs with kinase substrate consensus sequences. Substrates designed for CDK5 showed no phosphorylation by CDK2, opening the possibility for CDK5-specific assay development for selective drug discovery.

---

Since its discovery over 30 years ago^1^, cyclin-dependent kinase 5 (CDK5), a proline-directed serine/threonine protein kinase, has continually garnered attention largely due to its role in Alzheimer’s disease (AD)^2^ and other neurological disorders.^3^ More recently, CDK5 has also been implicated in cancer.^4^ Although CDK5 shares a close sequence homology with other cyclin-dependent kinases (CDKs)^5-7^, its activity is predominantly relegated to post-mitotic neurons where it is activated not by cyclins, but by the regulatory protein p35.^8^ p35-regulated CDK5 activity is essential for neuronal development, synaptic function, and cognitive processes such as learning and memory.^3^ Within cells, physiological CDK5/p35 activity is constrained to the periphery near the cellular membrane^9^ to which p35 is anchored by an N-terminal myristoyl moiety.^2,10^ However, upon exposure to neurotoxic signals, a cellular influx of Ca^2+^ activates proteolytic calpain to cleave p35, thereby releasing protein p25 from the membrane-bound p10.^11^ In addition to enabling CDK5 to access more potential substrates with its acquired cellular mobility, p25 has a half-life 10-fold longer than p35. This allows for the more stable and prolonged activation of CDK5 by p25, resulting in the dysregulation of CDK5 activity observed in neurodegenerative disorders.^2,12^ Dysregulated CDK5/p25 activity contributes to all three histopathological hallmarks of AD: extracellular deposition of β-amyloid (Aβ) plaques, formation of neurofibrillary tangles (NFT), and neuron death.^3^ Such hallmarks are the consequences of AD, but much less is known about the molecular interactions leading to this disease. The dysregulation of CDK5, a tau kinase, results in tau hyperphosphorylation and subsequent NFT formation.^13,14^ Aberrant CDK5/p25 activity has also been shown to induce Aβ accumulation through the upregulation of amyloid precursor protein (APP)^15^ and by activating beta-site amyloid precursor protein cleaving enzyme 1 (BACE1) both through direct phosphorylation^16^, and via the STAT3 pathway.^17^ CDK5/p25 triggers p53^18^ and JNK3^19^ pathways in addition to other apoptotic consequences by initiating nuclear envelope dispersion, DNA damage, mitochondrial dysfunction, while feeding forward additional oxidative stress and Ca^2+^ overload.^3,20^ Additional tools are needed to clearly understand the role aberrant CDK5 activity plays in AD and each of its hallmarks, and the effect CDK5 inhibition has on slowing and reversing AD progression.

Although hyperactive CDK5 has been clearly implicated in AD^13,21^, inhibitors which target CDK5 have yet to be approved for clinical treatment.^22^ This is largely due to the lack of selectivity of classified CDK5 inhibitors over other closely related CDKs such as CDK2, CDK5’s closest homolog, without incurring harmful, off-target effects.^23^ Just as no inhibitors for CDK5 have been approved, assays which specifically detect CDK5 activity among closely related kinases are limited, which makes screening inefficient. By developing synthetic peptide substrates capable of distinguishing the activities of these closely related kinases, we aim to provide improved assays as tools that could be used for selective kinase inhibitor drug discovery.

In recent years, the Parker Lab has developed tyrosine kinase assays featuring synthetic peptide substrates designed by combining kinase recognition sequences^24^ with various detection techniques including antibody-free time-resolved terbium (Tb) luminescence.^25^ Like homogenous time-resolved fluorescence (HTRF) assays, these assays are nonradioactive and well-suited for high-throughput inhibitor screening; however, the advantage is they do not require specific antibodies which are costly and often unavailable for phosphosites of interest, restricting such HTRF assays to the use of generic instead of kinase-specific substrates.^26^

The present work describes the creation of both LCMS and time-resolved terbium luminescence in vitro assays capable of detecting CDK5 activity over its closest relative CDK2 (62% sequence homology).^5^ This was accomplished by designing peptides that feature a previously reported consensus sequence for CDK5 (KHHKSPKHR)^27^ and implementing these peptides as assay substrates. To ensure C18 column retention for the LCMS assay, four leucine residues were placed on the N-terminus forming 4L-CDK5tide (Table 1). For the higher throughput terbium-based readout, versions of the substrate with a phospho-inducible Tb-chelating/sensitizing motif (as reported by the Zondlo Lab^28^) were also designed with the CDK5 substrate sequence. This enables an antibody- and fluorophore-free CDK5 assay featuring a synthetic substrate that luminesces upon phosphorylation. Specifically, this was achieved by merging the CDK5 consensus sequence with each of two Tb-sensitizing motifs reported by the Zondlo Lab, DKDADXWXS (Tb1) and DKDADXXWS (Tb2), in which X denotes any residue, resulting in Tb1-CDK5tide and Tb2-CDK5tide, respectively. By this design, excitation of tryptophan at 280 nm and energy transfer to a terbium ion (Tb^3+^), chelated nearby by phosphoserine and acidic residues of the Tb motif, leads to sensitized Tb luminescence (Fig. 1).

**Table 1:**
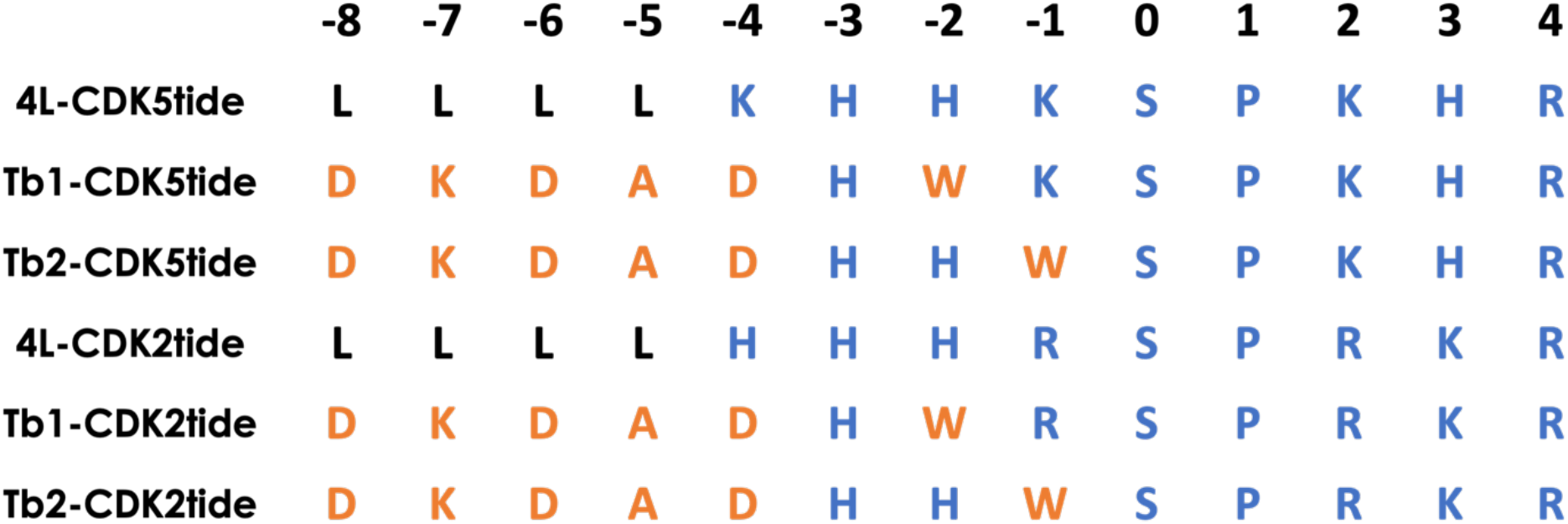
Substrates designed to assay CDK and CDK2. Residues DKDAD and W (orange) located N-terminal of S (phosphorylation site, position 0) are incorporated for phospho-inducible terbium chelation and as a UV acceptor, respectively.^28^ These residues are substituted into previously reported substrate consensus sequences of CDK5^27^ and CDK2^32^ (residues in blue). Peptides 4L-CDK5tide and 4L-CDK2tide are comprised of the corresponding kinase consensus sequence with N-terminal leucine residues (black) to ensure column retention for LCMS assay detection.

**Fig. 1:**
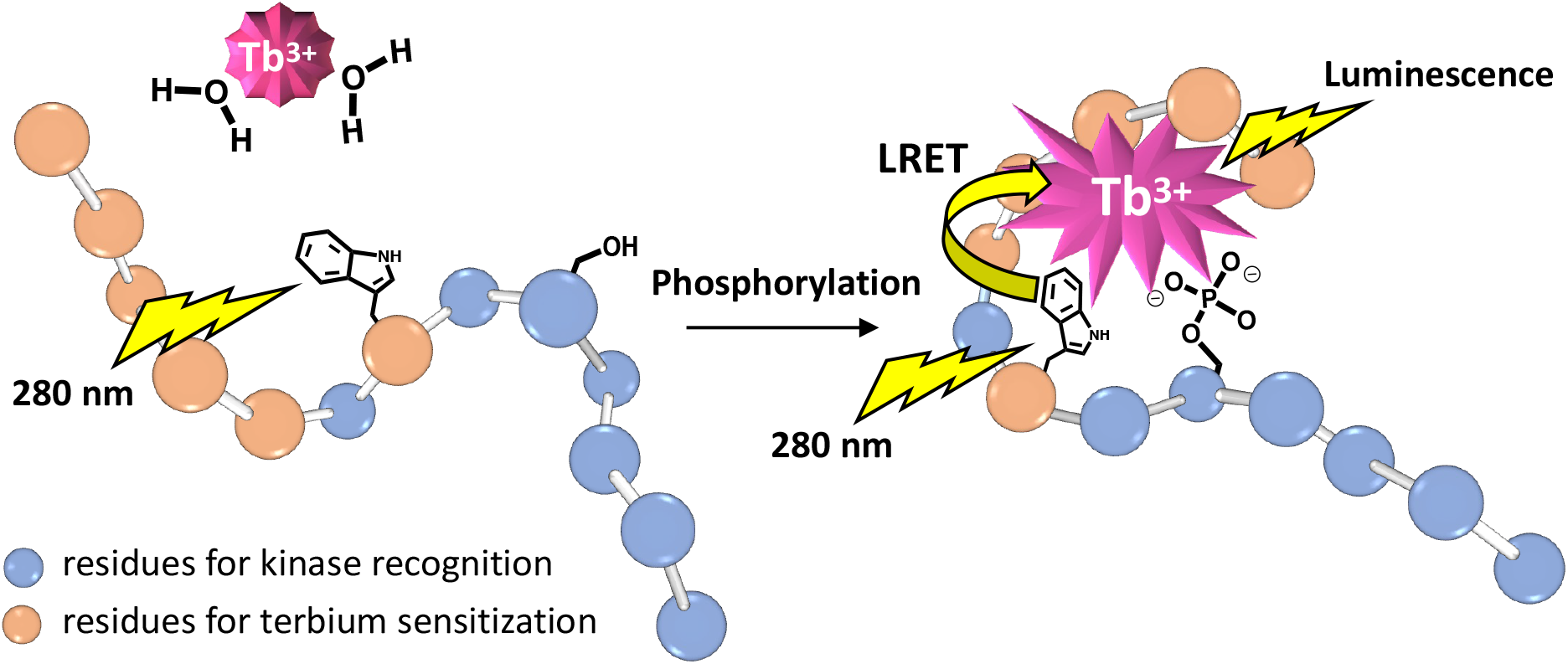
Time-resolved terbium luminescence assay featuring a synthetic peptide designed by merging a serine kinase consensus substrate sequence (blue residues) with a phospho-inducible terbium sensitization motif (orange residues). Upon kinase phosphorylation, the peptide chelates and sensitizes terbium. Excitation of a nearby positioned tryptophan residue (donor) at 280 nm and subsequent energy transfer to chelated terbium (acceptor) allow for luminescence.

While some background is observed for the unphosphorylated substrate, a calibration experiment with synthetic phosphopeptide standards of the Tb-sensitizing peptides demonstrated luminescence proportional to amount of phosphopeptide present (Fig. S13). Of the Tb-sensitizing used in this experiment, Tb1-CDK5tide displayed the greatest dynamic range between 100% phosphorylated vs. 100% non-phosphorylated (Fig. S13A, S13E). To show kinase activity with these substrates, CDK5/p25 kinase reactions were performed with 4L-CDK5tide, monitored with LCMS, and Tb1-CDK5tide and Tb2-CDK5tide, detected via luminescence. We confirmed that the peptides were indeed CDK5 substrates, and phosphorylation was easily monitored and quantified (Fig. 2A, solid symbols). Substrates Tb1-CDK5tide and Tb2-CDK5tide were phosphorylated less rapidly by CDK5 than 4L-CDK5tide, which was not unexpected since residues of the consensus sequence favourable to CDK5 were replaced at specific positions allowing for Tb chelation and sensitization. Nonetheless, phosphorylation of the Tb-CDK5tides was still efficient enough to be useful in assays, with the advantage that luminescence readouts can be performed in less than 30 seconds per sample. Of the two substrates designed for Tb luminescence, Tb1-CDK5tide with tryptophan at position -2 was phosphorylated only slightly more rapidly by CDK5 than Tb2-CDK5tide, which features tryptophan directly adjacent to the serine phosphorylation site (position -1). This suggests that the placement of the bulkier residue near the phosphorylation site was not a major issue, and the activity was more affected by replacing basic residues favoured by CDK5.

**Fig. 2:**
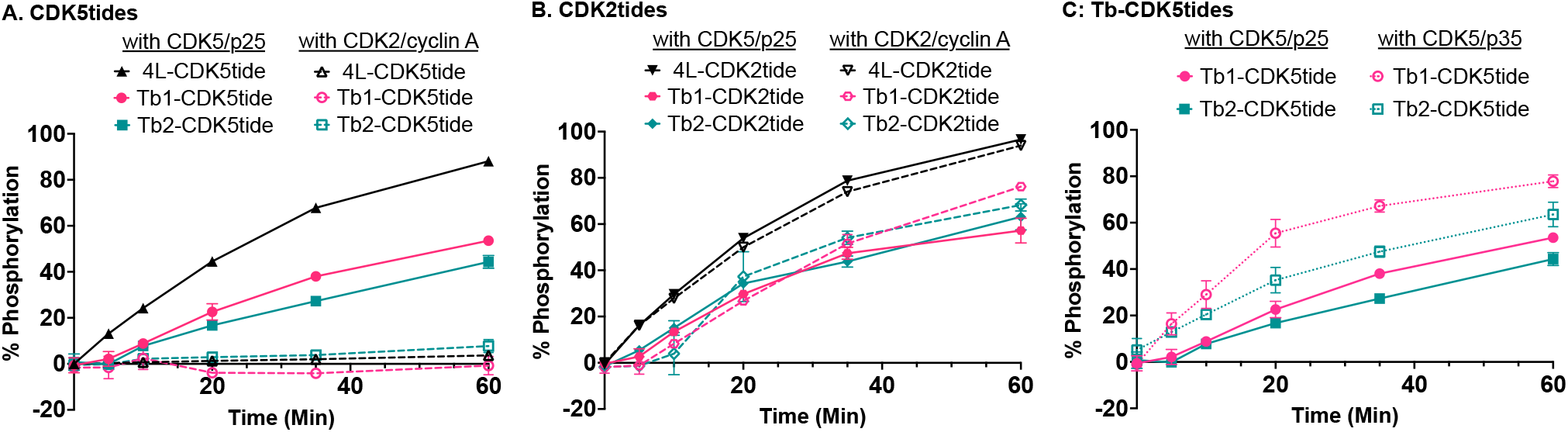
Reaction progress in percent phosphorylation of **A)** peptide substrates (CDK5tides) designed for CDK5 assayed with CDK5/p25 (solid symbols) and CDK2/cyclin A (open symbols); **B)** peptide substrates (CDK2tides) designed for CDK2 assayed with CDK5/p25 (solid symbols) and CDK2/cyclin A (open symbols); and **C)** Tb1-CDK5tide and Tb2-CDK5tide assayed with CDK5/p25 (solid symbols) and CDK5/p35 (dotted symbols). All reactions were performed in triplicate at 25°C using 25 µM substrate with 10 nM kinase/regulatory protein in kinase reaction buffer (25 mM HEPES, 50 µM ATP, 10 mM MgCl2, 0.2 mg/mL BSA, 0.2 mM DTT, pH 7.5). Reactions using 4L-CDK5tide were monitored by LCMS. Reactions using substrates Tb1-CDK5tide and Tb2-CDK5tide were monitored with time-resolved terbium luminescence assays.

Next, we tested whether these substrates could provide selective assays to identify inhibitors for CDK5 over CDK2. CDKs are known to share many similarities in structure^7^ and substrate preferences.^6,29^ Since CDK2 is structurally very similar to CDK5^30,31^, and reported substrate consensus sequences of CDK5^27^ and CDK2^32^ are also very similar^33,34^, we were surprised to find that all three substrates designed to assay CDK5 showed minimal to no product formation in the presence of CDK2/cyclin A (Fig. 2A, open symbols). To confirm the activity of CDK2/cyclin A, we applied a similar substrate design strategy to create a set of CDK2tides based on the CDK2 consensus sequence (Table 1).^32^ When treated with CDK2/cyclin A, each of the peptides 4L-CDK2tide, Tb1-CDK2tide and Tb2-CDK2tide were readily phosphorylated showing that our CDK2/cyclin A complex was indeed active (Fig. 2B, open symbols). However, unlike the CDK5tide substrates, which were phosphorylated only by CDK5, the CDK2tide substrates, were found to be phosphorylated by both CDK2 and CDK5 almost equally (Fig. 2B).

Encouraged by the ability of designed CDK5tides to selectively assay CDK5/p25 over its closest homolog CDK2, we were curious to test them with CDK5/p35, as substrate selectivity of CDKs is often directed by the interacting regulatory protein.^6,7,29^ When assayed with CDK5/p35, a higher rate of phosphorylation was observed for both Tb1-CDK5tide and Tb2-CDK5tide than in the CDK5/p25 assay performed under the same conditions (Fig. 2C). This was not completely unexpected since the consensus substrate sequence for CDK5 was obtained using CDK5/p35.^27^ Thus, the difference in rate of phosphorylation seen here between CDK5 when regulated by p35 vs. p25 could be due to either a shift in substrate preference (and relative activity on this particular substrate) guided by the regulatory protein or an intrinsic difference in activity between the two versions.

To better understand the selectivity of the CDK5 substrates, we examined the previously published crystal structure^30^ of CDK2/cyclin A3 complexed with the substrate HHASPRK (Fig. 3A) derived from the CDK2 substrate consensus sequence.^32^ Although CDK2 is the closest homolog of CDK5 with many similarities in sequence, structure, substrate preferences, and inhibitor profiles,^22,23^ they also have several key differences. In addition to not being involved in the cell cycle or regulated by a cyclin like CDK2, the mechanism of activation for CDK5 is different.^35,36^ CDK2 has a dual mechanism of activation^37,38^ that requires both the binding of a regulatory cyclin such as cyclin A or cyclin E^39^ and the phosphorylation of a threonine residue (T160) on its activation T-loop.^38,40^ The presence of phosphorylated T160 (pT160) on the activation loop of CDK2 drives a strong preference for lysine at position +3 of the substrate.^30^ The published crystal structure revealed the formation of a hydrogen bond between pT160 of active CDK2 and the +3-lysine residue of the substrate (Fig. 3A). Further, it has also been previously shown that substitution of the +3-lysine with alanine greatly reduced substrate phosphorylation by fully active CDK2.^41^ In the present work, the lack of phosphorylation of the CDK5tide substrates by CDK2 is most likely due to the placement of histidine instead of lysine in position +3 (Table 1). This suggests that CDK2 does not simply prefer a basic residue in the +3-substrate position, but that this residue should also possess a sidechain long enough to form a favourable interaction with pT160.

**Figure 3:**
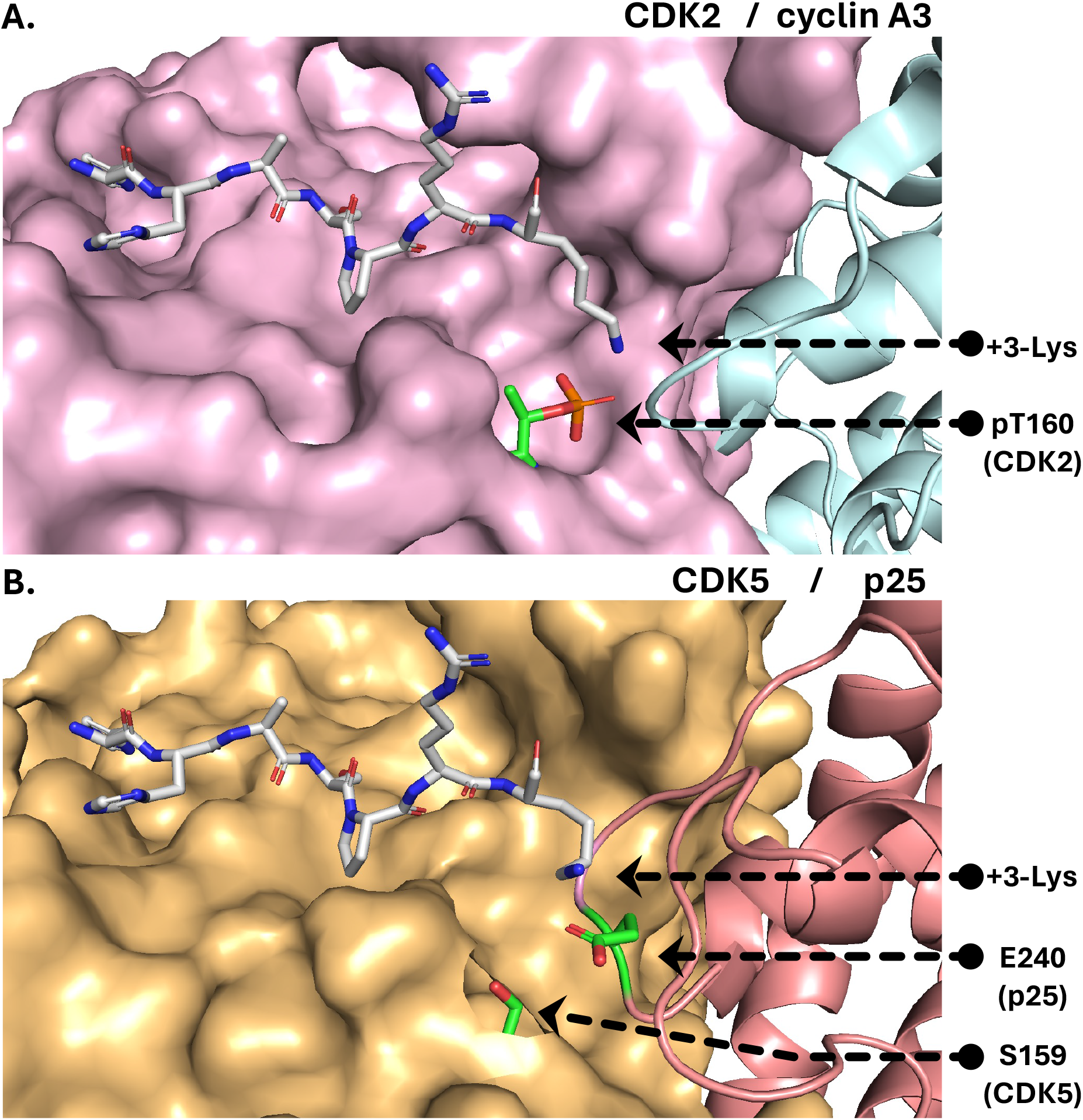
**A)** Crystal structure of substrate HHASPRK in the binding cleft of CDK2/cyclin A3 (PDB:1QMZ)^30^ and **B)** in-silico model of HHASPRK from PDB:1QMZ (with +3-lysine rotamer) added and aligned with crystal structure of CDK5/p25 (PDB: 1H4L)^31^. Labels added highlighting interaction of +3-lysine with pT160 of CDK2 (A) and E240 of p25 (B). Figures generated in Pymol from published structural models.

CDK5 on the other hand only requires regulatory protein binding to become active. Although CDK5 possesses a serine residue at position 159 (S159), analogous to T160 in CDK2, it does not require phosphorylation to become active.^42^ Furthermore, structural studies suggest that phosphorylation of S159 is sterically disfavoured and substitution of S159 with glutamate (as an analogue of phosphoserine) impaired p35 binding and greatly reduced activity.^31^ In terms of structure, p35 and p25 form more contacts with CDK5 than cyclin A with CDK2, thus the mere binding of the regulatory protein is enough to tether the CDK5 activation loop forming an extended conformation resembling that of CDK2 dually activated by phosphorylation and cyclin binding (Fig. 3B).^30,31,35^ In addition to forming more contacts than cyclin A with the respective kinase, p35/p25 has been shown to interact directly with the substrate. Tarricone et al. showed that a glutamate residue located in position 240 (E240) of p35/p25 imparts substrate selectivity for basic residues at position +3 in a manner similar to pT160 of CDK2 (Fig. 3B).^31^ Upon substitution of E240 with alanine and glutamine, activity was greatly decreased and the ability to distinguish between lysine and alanine in the +3-substrate position was lost. This suggests that even without phosphorylation of activation loop S159, CDK5 maintains a strong preference for a basic residue located in substrate position +3, and helps explain its ability to phosphorylate both CDK5tides (+3-histidine) and CDK2tides (+3-lysine) in our experiments in contrast to CDK2.

In this work, a set of synthetic peptide substrates were developed that can be used to specifically assay CDK5 over its close homolog CDK2. This set includes 4L-CDK5tide designed for LCMS detection from the CDK5 substrate consensus sequence, as well as substrates Tb1-CDK5tide and Tb2-CDK5tide, which merge Tb-sensitizing motifs with the consensus sequence. Incorporation of Tb-sensitizing motif allows for antibody-free, time-resolved luminescence proportional to phosphopeptide produced that will be compatible with high-throughput assay detection. These substrates could be used for multiplexed dual screening of CDK5 and CDK2, and may also be useful for detecting CDK5 vs. CDK2 activity in biological samples.

## Notes and references

We acknowledge and appreciate funding support for this work from National Institutes of Health (NIH) grants R01CA182543, R01GM146386 and T32AG029796.

