## Supplementary Information for "Antibody-free time-resolved terbium luminescence assays designed for cyclin-dependent kinase 5 (CDK5)"

**Table of Contents** **Page**

**Materials and Methods:** 2

Peptide Synthesis: 2

Peptide Characterisation by LCMS 2

**Figure S1. 4L-CDK5tide:** $H_2N$-LLLLKHHKSPKHR-$CONH_2$ 2

**Figure S2. 4L-CDK5tide (phosphorylated):** $H_2N$-LLLLKHHKpSPKHR-$CONH_2$ 3

**Figure S3. Tb1-CDK5tide:** $H_2N$-DKDADHWKSPKHR-$CONH_2$ 3

**Figure S4. Tb1-CDK5tide (phosphorylated):** $H_2N$-DKDADHWKpSPKHR-$CONH_2$ 3

**Figure S5. Tb2-CDK5tide:** $H_2N$-DKDADHHWSPKHR-$CONH_2$ 4

**Figure S6. Tb2-CDK5tide (phosphorylated):** $H_2N$-DKDADHHWpSPKHR-$CONH_2$ 4

**Figure S7. 4L-CDK2tide:** $H_2N$-LLLLHHHRSPRKR-$CONH_2$ 4

**Figure S8. 4L-CDK2tide (phosphorylated):** $H_2N$-LLLLHHHRpSPRKR-$CONH_2$ 5

**Figure S9. Tb1-CDK2tide:** $H_2N$-DKDADHWRSPRKR-$CONH_2$ 5

**Figure S10. Tb1-CDK2tide (phosphorylated):** $H_2N$-DKDADHWRpSPRKR-$CONH_2$ 5

**Figure S11. Tb2-CDK2tide:** $H_2N$-DKDADHHWSPRKR-$CONH_2$ 6

**Figure S12. Tb2-CDK2tide (phosphorylated):** $H_2N$-DKDADHHWpSPRKR-$CONH_2$. 6

Kinase Assays: 6

**Figure S13. Terbium Luminescence Assay Calibration** 7

**Materials and Methods:**

Peptide Synthesis:

Peptide substrates were synthesized using standard Fmoc chemistry on Rink amide resin (0.05 mmol, Gyros Protein Technologies) with a Symphony X automated peptide synthesizer (Gyros Protein Technologies). Fmoc deprotection was performed in 20% piperidine (2 x 5 min) and peptide elongation was carried out by coupling Fmoc-protected amino acids (6 equiv., Gyros Protein Technologies) applying HCTU (5.7 equiv., Gyros Protein Technologies) and NMM (12 equiv., Gyros Protein Technologies) for activation (2 x 20 min). The coupling of phosphoserine (Fmoc-Ser(PO(OBzl)OH)-OH, 4 equiv., BIOSYNTH) was achieved with 3.8 equiv. HCTU and 8 equiv. NMM (1 x 8h). The coupling times of three residues incorporated after the phosphoserine were extended from 2 x 20 min to 2 x 2h. Peptides were cleaved from resin, and their sidechains were deprotected in a cocktail of 94% trifluoroacetic acid (Sigma-Aldrich), 2.5% water, 2.5% ethanedithiol (Sigma-Aldrich) and 1% triisopropylsilane (ChemImpex). Following cleavage, peptides were precipitated and washed thrice with ice-cold diethyl ether (Fisher) before redissolving the final peptide pellet with aqueous acetonitrile and freeze drying.

Peptides were purified via RP-LCMS using an Agilent 1200 Series HPLC on an Agilent Zorbax 300SB-C18 column (5 micron, 9.4 x 250 mm) at flowrate of 4 mL/min. Eluted peptides were identified with an Agilent 6130A mass spectrometer. Prior to assay application, all peptides were analyzed via RP-LCMS using an Agilent 1200 Series HPLC connected to an Agilent 6130A MS on an Agilent Zorbax-C18 column (5 micron, 2.1 x 250mm) with a flowrate of 0.25 mL/min using a gradient of aqueous acetonitrile (with 0.1% formic acid) increasing from 0% acetonitrile at minute 10 to 30% at minute 30. Chromatographs and mass spectra for each of the peptides are shown below in figures S1-S12.

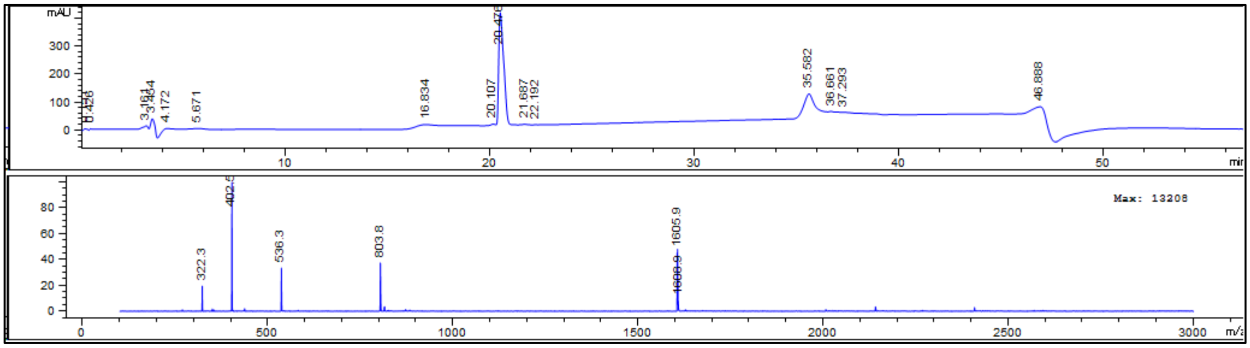

**Figure S1. 4L-CDK5tide:** $H_2N$-LLLLKHHKSPKHR-$CONH_2$; Mass Calculated: 1604.9; Found:1604.9

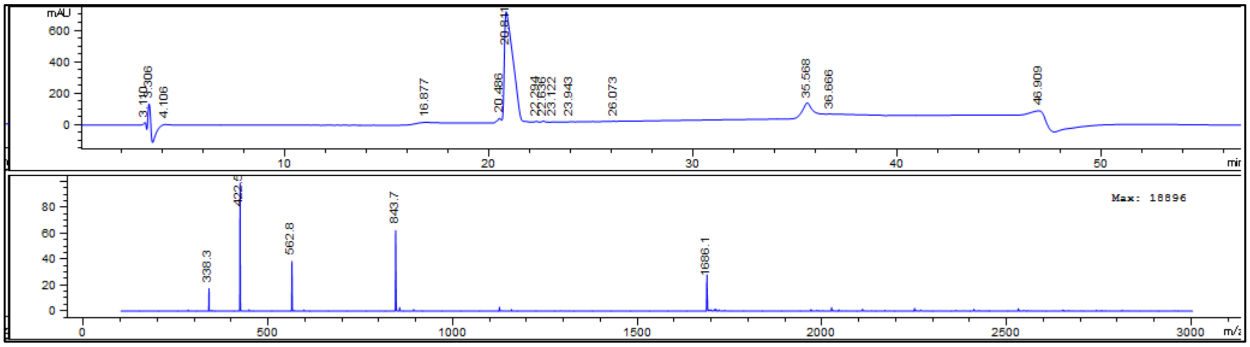


**Figure S2. 4L-CDK5tide (phosphorylated):** $H_2N$-LLLLKHHKpSPKHR-$CONH_2$; Mass Calculated: 1684.9; Found:1685.1

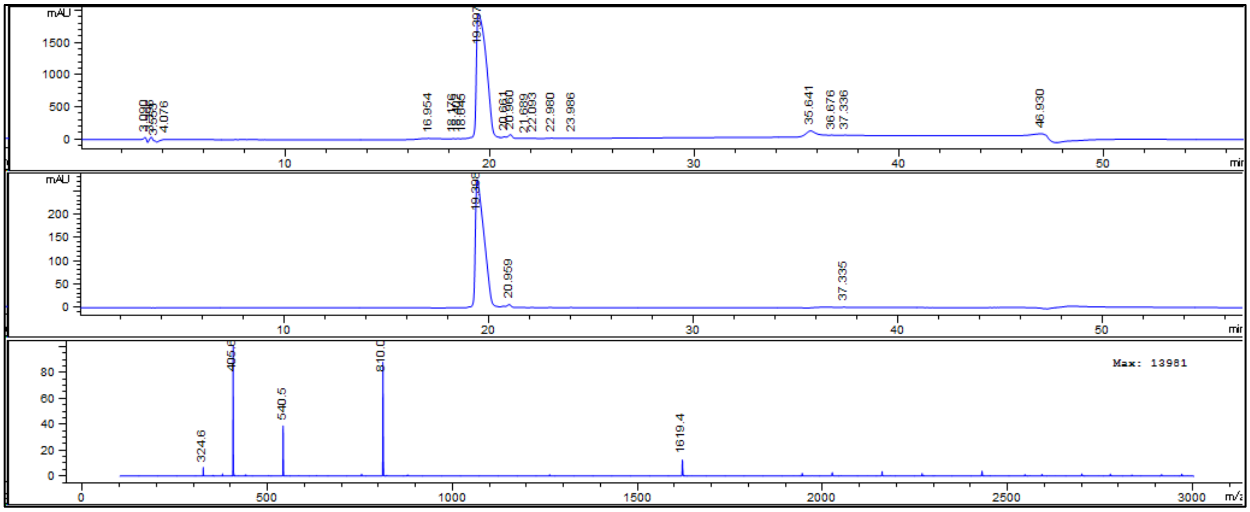


**Figure S3. Tb1-CDK5tide:** $H_2N$-DKDADHWKSPKHR-$CONH_2$; Mass Calculated: 1617.8; Found:1618.4

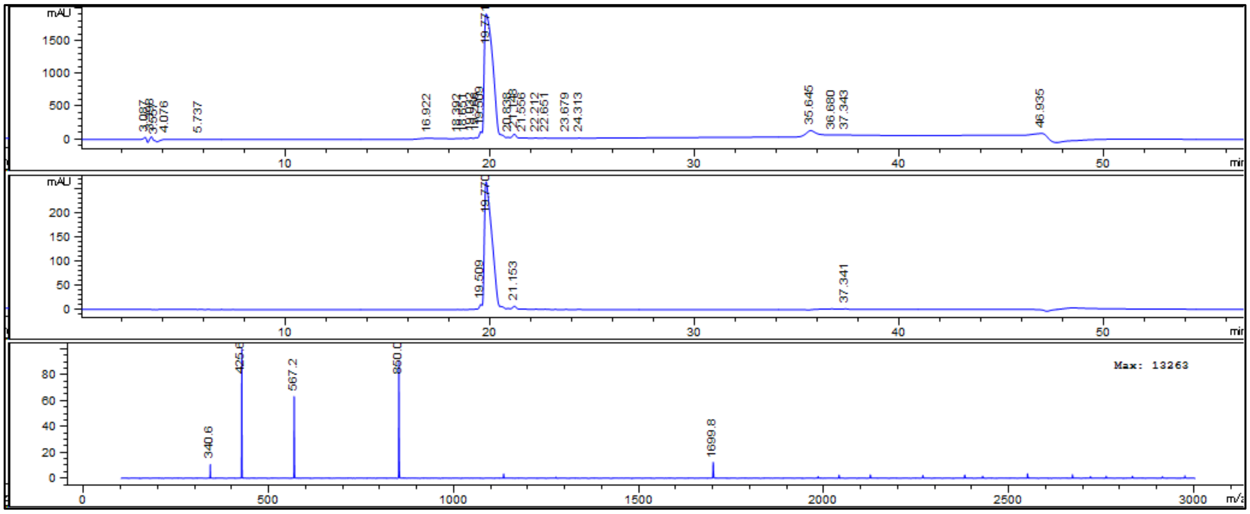


**Figure S4. Tb1-CDK5tide (phosphorylated):** $H_2N$-DKDADHWKpSPKHR-$CONH_2$; Mass Calculated: 1697.8; Found:1698.8

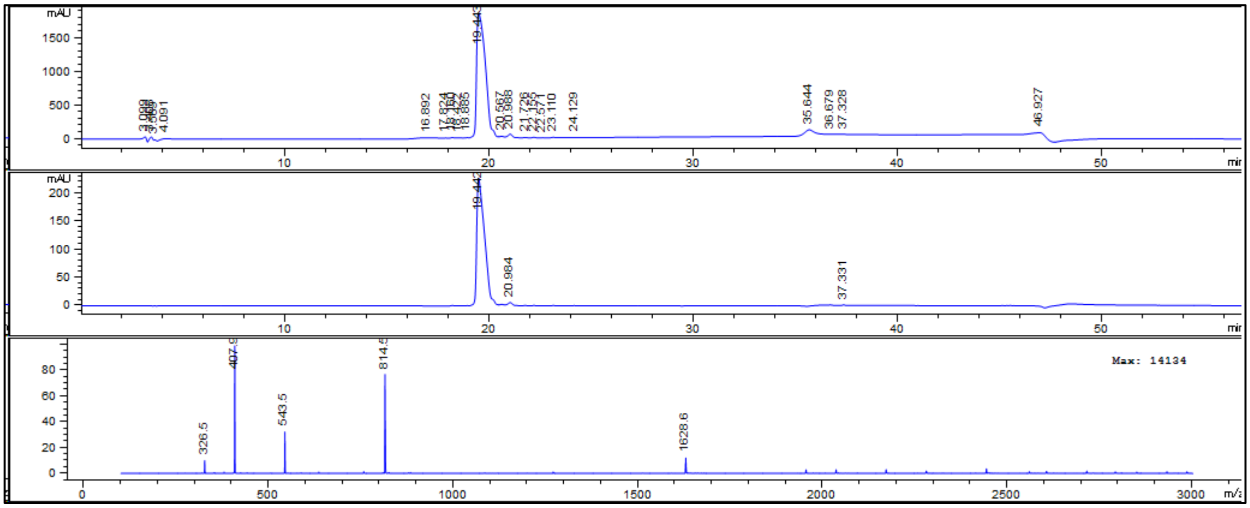


**Figure S5. Tb2-CDK5tide:** $H_2N$-DKDADHHWSPKHR-$CONH_2$; Mass Calculated: 1626.7; Found:1627.6

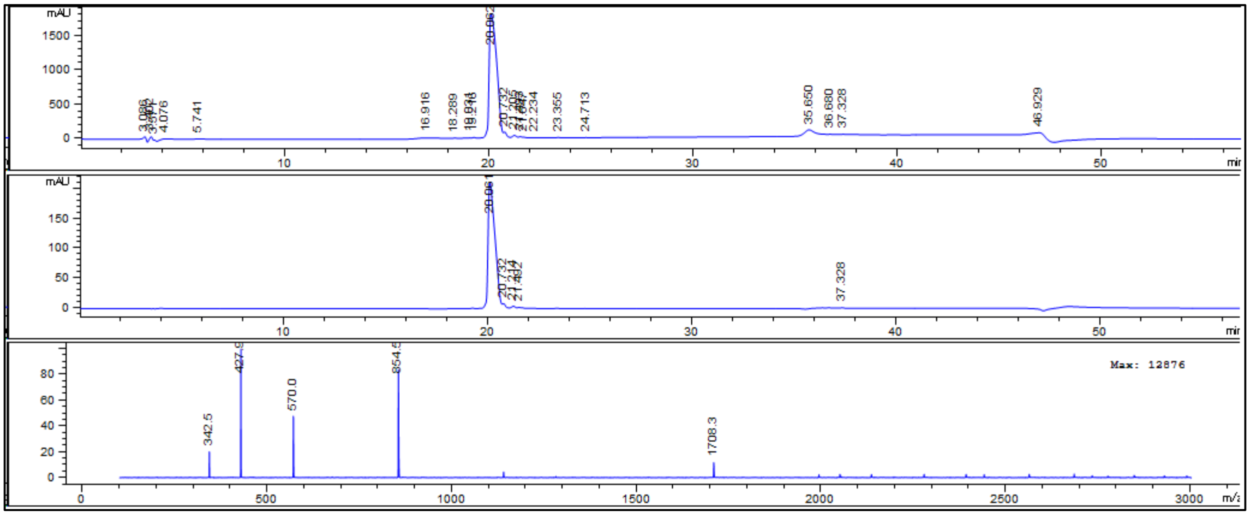


**Figure S6. Tb2-CDK5tide (phosphorylated):** $H_2N$-DKDADHHWpSPKHR-$CONH_2$; Mass Calculated: 1706.7; Found:1707.3

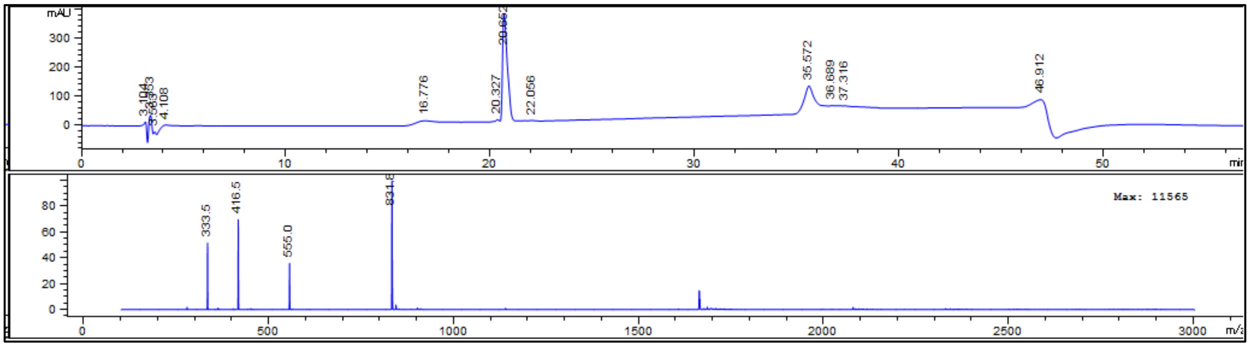


**Figure S7. 4L-CDK2tide:** $H_2N$-LLLLHHHRSPRKR-$CONH_2$; Mass Calculated: 1661.0; Found:1661.6

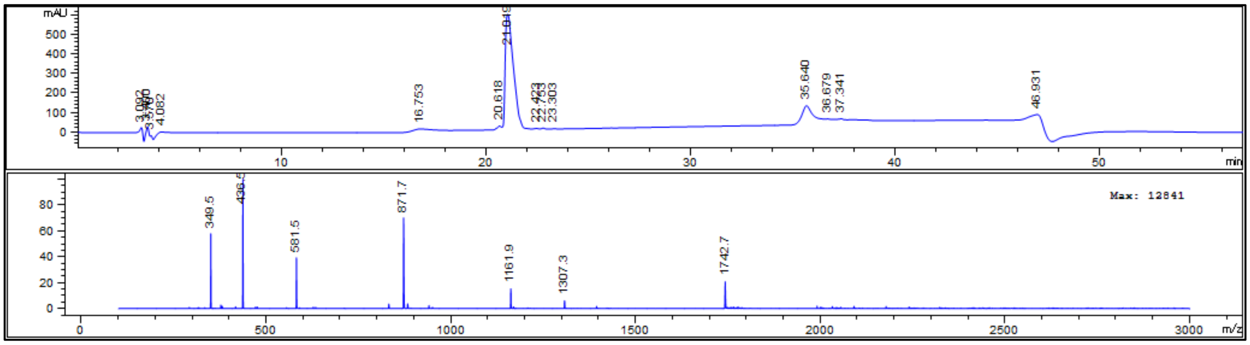


**Figure S8. 4L-CDK2tide (phosphorylated):** $H_2N$-LLLLHHHRpSPRKR-$CONH_2$; Mass Calculated: 1741.0; Found:1741.7

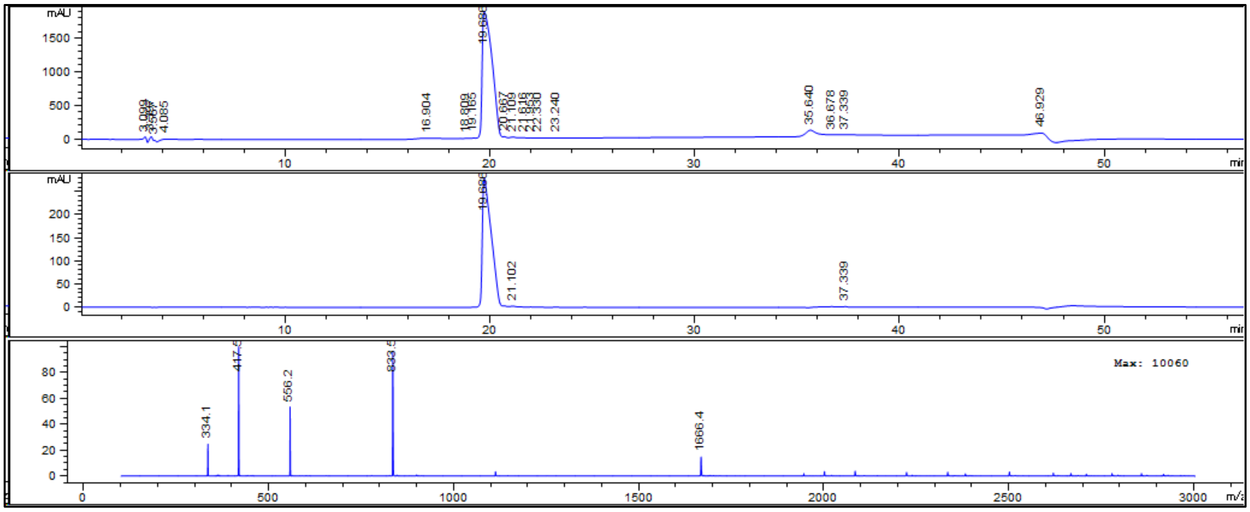


**Figure S9. Tb1-CDK2tide:** $H_2N$-DKDADHWRSPRKR-$CONH_2$; Mass Calculated: 1664.8; Found:1665.4

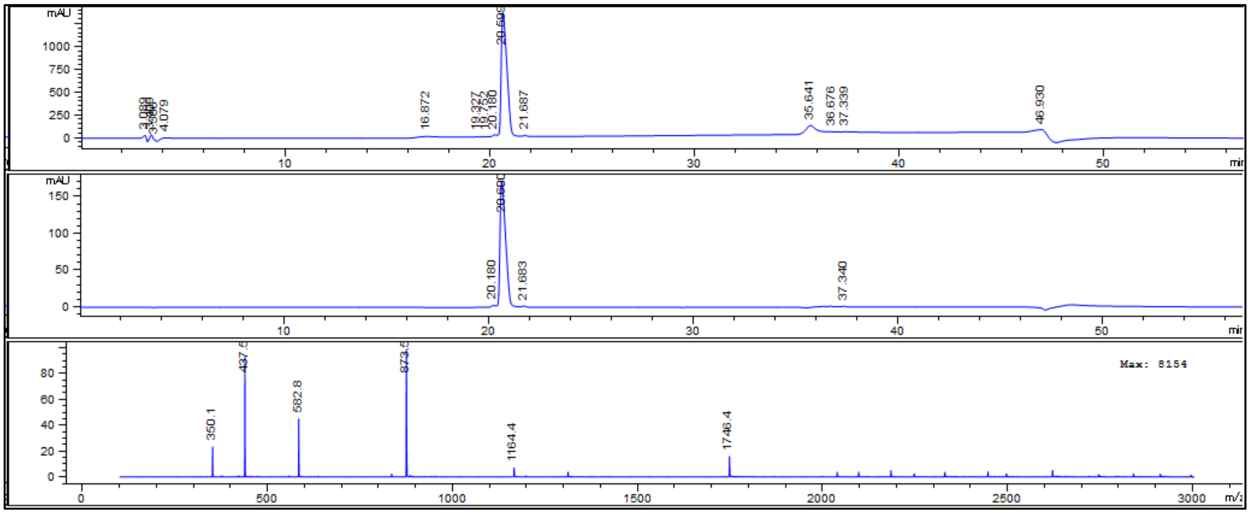


**Figure S10. Tb1-CDK2tide (phosphorylated):** $H_2N$-DKDADHWRpSPRKR-$CONH_2$; Mass Calculated: 1744.8; Found:1745.4

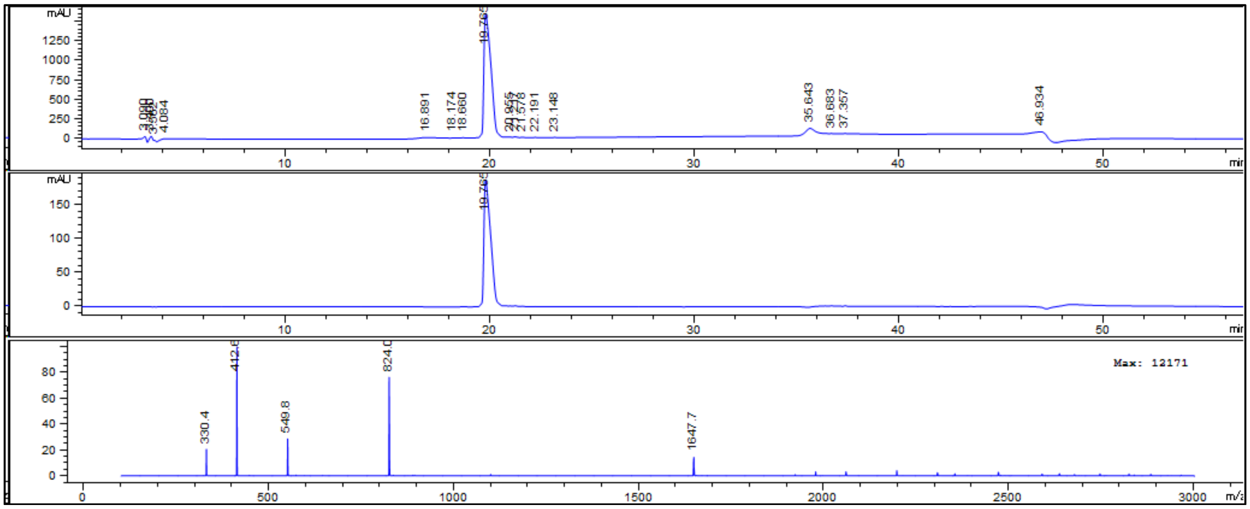


**Figure S11. Tb2-CDK2tide:** $H_2N$-DKDADHHWSPRKR-$CONH_2$; Mass Calculated: 1645.8; Found:1646.7

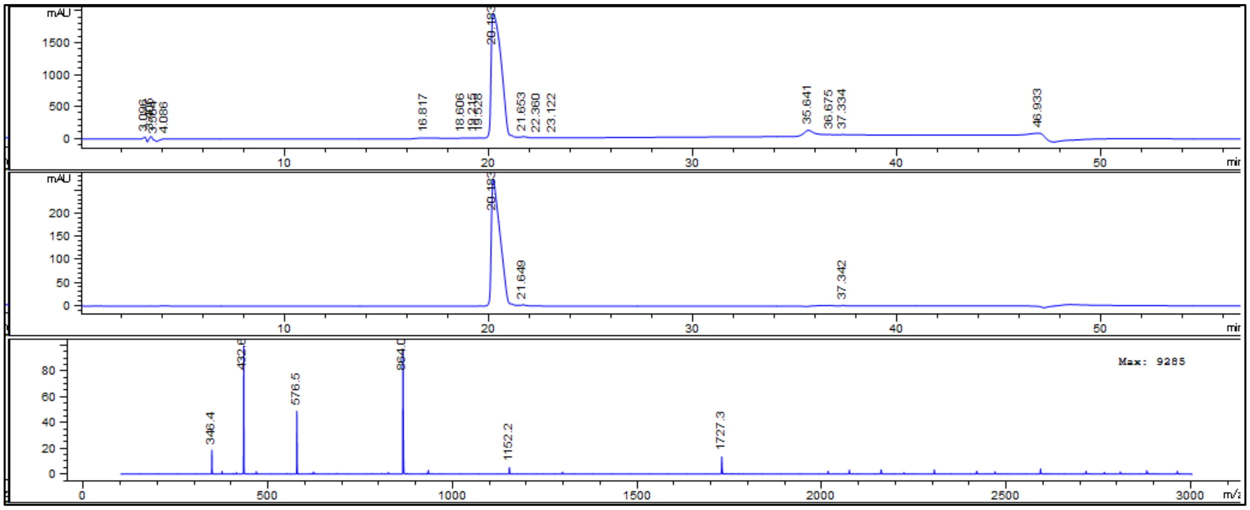


**Figure S12. Tb2-CDK2tide (phosphorylated):** $H_2N$-DKDADHHWpSPRKR-$CONH_2$; Mass Calculated: 1725.8; Found:1726.3

Kinase Assays:

Kinase assays were performed in triplicate at 25°C with 10 nM kinase/regulatory protein (SignalChem) in kinase reaction buffer (25 mM HEPES, 50 µM ATP, 10 mM $MgCl_2$, 0.2 mg/mL BSA, 0.2 mM DTT, pH 7.5). Kinase reactions were initiated with the addition of substrate to achieve an initial reaction concentration of 25 µM. Reaction aliquots (45 µL) were withdrawn and quenched at 5, 10, 20, 35 and 60 minutes.

For assays detected by HPLC, reaction aliquots were quenched upon addition to 15 µL aqueous TFA (20%). Quenched aliquots (50 µL) were injected and run at 0.25 mL/min on HPLC (Agilent 1200 series) using an Agilent Zorbax-C18 column (5 micron, 2.1 x 250mm) from 3% to 10% aqueous acetonitrile with 0.1% formic acid over 20 minutes to fully separate products from substrates. Identities of substrates and products were further confirmed with MS (Agilent 6130A). Reaction progress was monitored by detecting the absorbance at 214 nm (Agilent 1100 Series G1315B) of eluted substrates and products present at each timepoint and integrating the corresponding peaks. These values were normalized by dividing the area corresponding to the product's peak by the summed areas of both the product and the substrate. The normalized values were further calibrated using standard curves established for each reaction using peptide solutions containing known ratios of product phosphopeptide and substrate in buffer and quenching conditions mimicking the reaction aliquots, however, without kinase. Since phosphorylation has a minimal effect on absorbance at 214 nm, changes imparted by calibration were minor (>2%).

For assays detected by terbium luminescence, aliquots were quenched in 45 µL 6 M urea. The quenched aliquots were treated with 22.5 µL of detection solution (0.5 mM $TbCl_3$, 0.5 M NaCl) and added (100 µL) to a 384-well black flat-bottom plates (PP-MICROPLATE, Greiner bio-one). Time-resolved terbium luminescence emission was measured using a BioTek Synergy Neo 2 plate reader. Emission spectra were collected between 450 and 650 nm (10 nm bandwidth, 2 nm step) with 50 µs delay following excitation at 280 nm (10 nm bandwidth). Readings (20/datapoint) were taken for 1 ms at a read height of 7 mm with the gain adjustment at 180, and the plate reader lamp energy set to high sensitivity. Luminescence was quantified by integrating the terbium emission spectra. The integrated luminescence signals corresponding to each of the reaction timepoints were converted to % phosphorylation with calibration curves (Fig. S13). Calibration curves were established for each reaction using standard peptide solutions containing known ratios of product phosphopeptide and substrate (10 µM total peptide concentration) in buffer and quenching/detection conditions mimicking the reaction aliquots, however, with the exclusion of kinase (Fig. S13A-D).

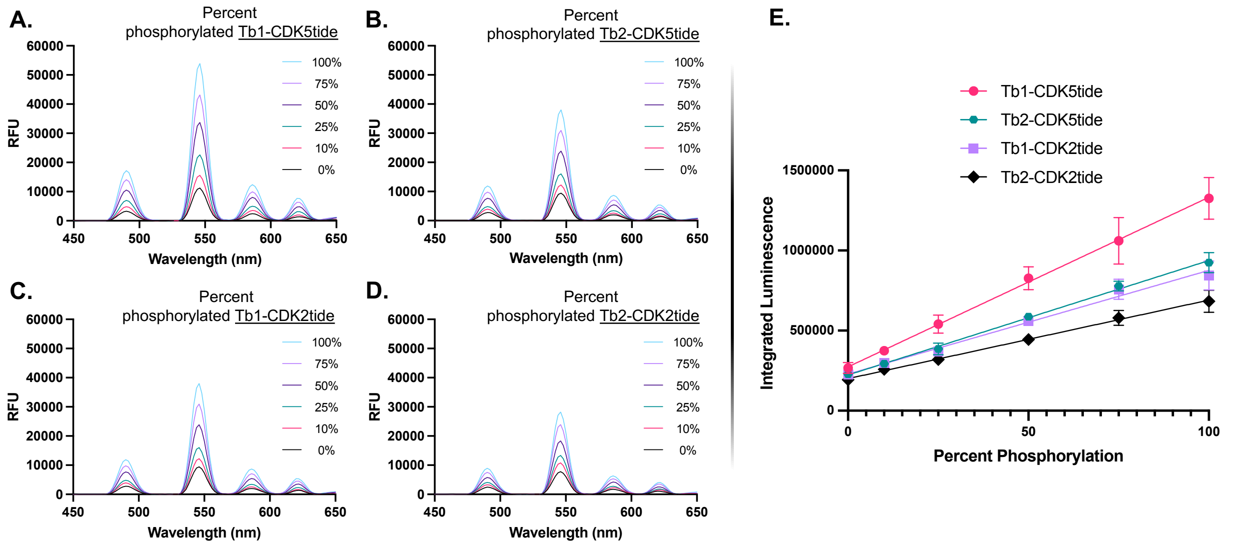


**Figure S13. Terbium Luminescence Assay Calibration:** Luminescence emission spectra in relative fluorescence units (RFU) corresponding to standard peptide solutions containing known ratios of phosphopeptides (kinase products) with their respective substrates **A)** Tb1-CDK5tide, **B)** Tb2-CDK5tide, **C)** Tb1-CDK2tide, **D)** Tb2-CDK2tide, and **E)** resulting calibration curves used to convert integrated emission spectra to reaction progress in percent phosphorylation for each substrate reacted with kinases CDK5/p25 and CDK2/cyclin A. Standards contained 10 µM total peptide concentration in buffer and quenching/detection conditions (10 mM HEPES pH 7.5, 20 µM ATP, 4 mM $MgCl_2$, 80 µM DTT, 80 ng/µL BSA, 2.4 M Urea, 100 µM $TbCl_3$, 100 mM NaCl) mimicking quenched reaction aliquots.
